# Characterization of the adaptation to visuomotor rotations in the muscle synergies space

**DOI:** 10.1101/2019.12.12.873802

**Authors:** Giacomo Severini, Magdalena Zych

## Abstract

The adaptation to visuomotor rotations is one of the most studied paradigms of motor learning. Previous literature has presented evidence of a dependency between the process of adaptation to visuomotor rotations and the constrains dictated by the workspace of the biological actuators, the muscles, and their co-activation strategies, modeled using muscle synergies analysis. To better understand this relationship, we asked a sample of healthy individuals (N =7) to perform two experiments aiming at characterizing the adaptation to visuomotor rotations in terms of rotations of the activation space of the muscle synergies during isometric reaching tasks. In both experiments, subjects were asked to adapt to visual rotations altering the position mapping between the force exerted on a fixed manipulandum and the movement of a cursor on a screen. In the first experiment subjects adapted to three different visuomotor rotation angles (30°, 40° and 50° clockwise) applied to the whole experimental workspace. In the second experiment subjects adapted to a single visuomotor rotation angle (45° clockwise) applied to eight different sub-spaces of the whole workspace, while also performing movements in the rest of the unperturbed workspace. The results from the first experiment confirmed the observation that visuomotor rotations induce rotations in the synergies activation workspace that are proportional to the visuomotor rotation angle. The results from the second experiment showed that rotations affecting limited sub-spaces of the whole workspace are adapted for by rotating only the synergies involved in the movement, with an angle proportional to the distance between the preferred angle of the synergy and the sub-space covered by the rotation. Moreover, we show that the activation of a synergy is only rotated when the sub-space covered by the visual perturbation is applied at the boundaries of workspace of the synergy. We found these results to be consistent across subjects, synergies and sub-spaces. Moreover, we found a correlation between synergies and muscle rotations further confirming that the adaptation process can be well described, at the neuromuscular level, using the muscle synergies model. These results provide information on how visuomotor rotations can be used to induce a desired neuromuscular response.

## Introduction

Adaptation to visuomotor rotations is one of the most widely studied paradigms of motor learning (Krakauer et al., 2000; Krakauer et al., 2019), and has been extensively discussed in the past three decades. Correlates of the processes contributing to visuomotor adaptations have been observed, directly or indirectly, in the primary motor cortex (Wise et al., 1998), the supplementary motor cortex (Mandelblat-Cerf et al., 2009), the premotor cortex (Perich et al., 2018) and the cerebellum (Della-Maggiore et al., 2009; Schlerf et al., 2012; Block and Celnik, 2013), in both humans and animal models.

Despite these neurophysiological insights, most of what we know regarding the functional processes contributing to visuomotor adaptation has been obtained through behavioral experiments (Krakauer et al., 1999; Krakauer et al., 2000; Bock et al., 2001; Krakauer et al., 2006; Hinder et al., 2007; Brayanov et al., 2012; De Marchis et al., 2018). These experiments have allowed to characterize adaptations, and, consequently, the control of voluntary movements, from several different points of view. Some studies have characterized how adaptations generalize (Shadmehr, 2004), either by transferring to similar untrained scenarios (Krakauer et al., 2006), or even to another limb (Sainburg and Wang, 2002) or by interfering with incompatible adaptations (Bock et al., 2001; Woolley et al., 2007). Other studies have been able to discern between the implicit and explicit components of the learning associated with the adaptation process (Taylor et al., 2014; Bond and Taylor, 2015). Moreover, the visuomotor adaptation paradigm has often been used to investigate which frame of reference, implicit (joint-based) or explicit (world-based) is employed when planning, executing and adapting movements (Krakauer et al., 2000; Brayanov et al., 2012; Carroll et al., 2014; Rotella et al., 2015). Most of these studies have investigated adaptations in terms of task performance or through their unraveling in the intrinsic space of joint coordinates or in the extrinsic space specific to the experimental set-up that was employed in the study.

A few studies have also investigated how motor adaptations are achieved in the space of the body actuators, the muscles. In these studies, visuomotor and force-field adaptations have been linked to the “tuning” of muscular activity (Thoroughman and Shadmehr, 1999; Gentner et al., 2013), consisting in perturbation-dependent rotations of the activation workspace of the muscles involved in the movement. Following the observation that complex movements can be described, at the neuromuscular level, by the combination of a limited number of muscular co-activation modules, generally referred-to as muscle synergies (d’Avella et al., 2003; d’Avella et al., 2006; Delis et al., 2014), a number of studies have also attempted to characterize motor adaptations in relationship to the muscle synergies structure (de Rugy et al., 2009; Berger et al., 2013; Gentner et al., 2013; De Marchis et al., 2018). Such studies presented mounting evidence that the underlying structure of neuromechanical control directly constraints the adaptation process (de Rugy et al., 2009), correlates with phenomena such as generalization (De Marchis et al., 2018) and even appears to dictate what kind of perturbations can be adapted for (Berger et al., 2013). Nevertheless, a full characterization of the link between motor adaptations and the tuning of the muscle synergies is still lacking.

Therefore, the aim of this study is to further understand how the muscular co-activation strategies that have been observed consistently during voluntary movements in the upper limb constraint visuomotor adaptations and if there are identifiable and exploitable relationships between the spatial characteristics of a perturbing visuomotor rotation and the muscular activity during isometric reaching tasks.

To achieve these aims, we first investigated how different visuomotor rotation angles applied to the whole workspace during isometric reaching movements affect the rotation of all the synergies characterizing the neuromuscular control. The aim of this experiment was to confirm previous observations, derived in studies employing only one perturbation angle, that synergies and muscles tuning is proportional to the angle of the perturbing rotation (Gentner et al., 2013; De Marchis et al., 2018). In a second experiment we investigated how a rotation affecting a small sub-space of the whole movement workspace leads to differential rotations of the synergies involved.

Here we found a selective tuning of the muscle synergies that is constrained, as expected, only to the synergies directly acting in the perturbed sub-space and that is proportional to the distance between the perturbed workspace and the workspace covered by each synergy. This proportionality allowed us to derive some generalizable observations on how synergies and muscles are tuned in response to specific visuomotor rotations. The results of this study can provide useful information on how visuomotor rotations can be used to design a desired neuromuscular output, by exploiting fixed relationships between the representation of movement in the neuromuscular space and the visual perturbations.

## Methods

### Experimental setup and Protocol

Seven healthy individuals (2 females, age 26.7 ± 2.6) participated in this study. Each individual participated in two experimental sessions, performed in different days within the same week, each consisting of a series of isometric reaching tasks performed with their right arm. All the experimental procedures describe in the following have been approved by the Ethical Committee of University College Dublin and have been conducted according to the WMA’s declaration of Helsinki. All subjects gave written informed consent before participating to this study. Each experimental session was performed using the setup previously used in (De Marchis et al., 2018). During all experimental procedures, the subjects sat in a chair with their back straight and their right hand strapped to a fixed manipulandum. Their right forearm was put on a support plan, their elbow was kept flexed at 90° and their shoulder horizontally abducted at 45° (**Figure 1A**), so that the manipulandum would be exactly in front of the center of rotation of their shoulder. The wrist and forearm were wrapped to the support plan and immobilized using self-adhesive tape. The elevation of the chair was controlled so to keep the shoulder abducted at 100°. The manipulandum consisted of a metal cylinder of 4 cm of diameter attached to a tri-axial load cell (3A120, Interface, UK). Data from the load cell were sampled at 50 Hz. Subjects sat in front of a screen displaying a virtual scene at a distance of 1 m. The virtual scene consisted of a cursor, whose position was commanded in real-time by the *x* and *y* components of the force exerted on the load cell through the manipulandum, a filled circle indicating the center of the exercise space and, depending on the phase of the exercise, a target, represented by a hollow circle. Both the central and target circles had a radius of 1.3 cm. Across all the blocks of the experiment subjects experienced a total of 16 different targets, positioned in a compass-like configuration at angular distances of 22.5° (**Figure 1A**) and at a distance of 9.5 cm from the center of the screen, equivalent to 15 N of force exerted on the fixed manipulandum (with the center of the virtual scene corresponding to 0 N). The virtual scene and the exercise protocol were controlled using a custom Labview software. In both experiments, the subjects were asked to perform both unperturbed and perturbed movements, where the perturbation consisted of a clockwise visuomotor rotation affecting the mapping between the force exerted on the manipulandum and the position of the cursor shown on the virtual scene. The angle of the visuomotor rotation varied across the different experiments (see below). At the beginning of each experimental session subjects underwent a practice trial with the setup. In this trial (identical to the unperturbed baseline and post-adaptation trials present in both Experiment 1 and 2), subjects were asked to reach to the 16 targets in a randomized order three times, for a total of 48 movements. In all the trials the movement time was not restricted, and subjects were presented a new target only when the current target had been reached. However, subjects were instructed to reach the targets at a comfortable speed in a time not exceeding 1.5 s and were given negative feedback (target turning red) if they took more than the expected time to reach for each target. Subjects were asked to bring the cursor back to the center of the screen as soon as they reached a target. These instructions were used for all perturbed and unperturbed reaching trials performed during both experiments, with the exclusion of the normalization blocks (see below).

**Figure 1.**
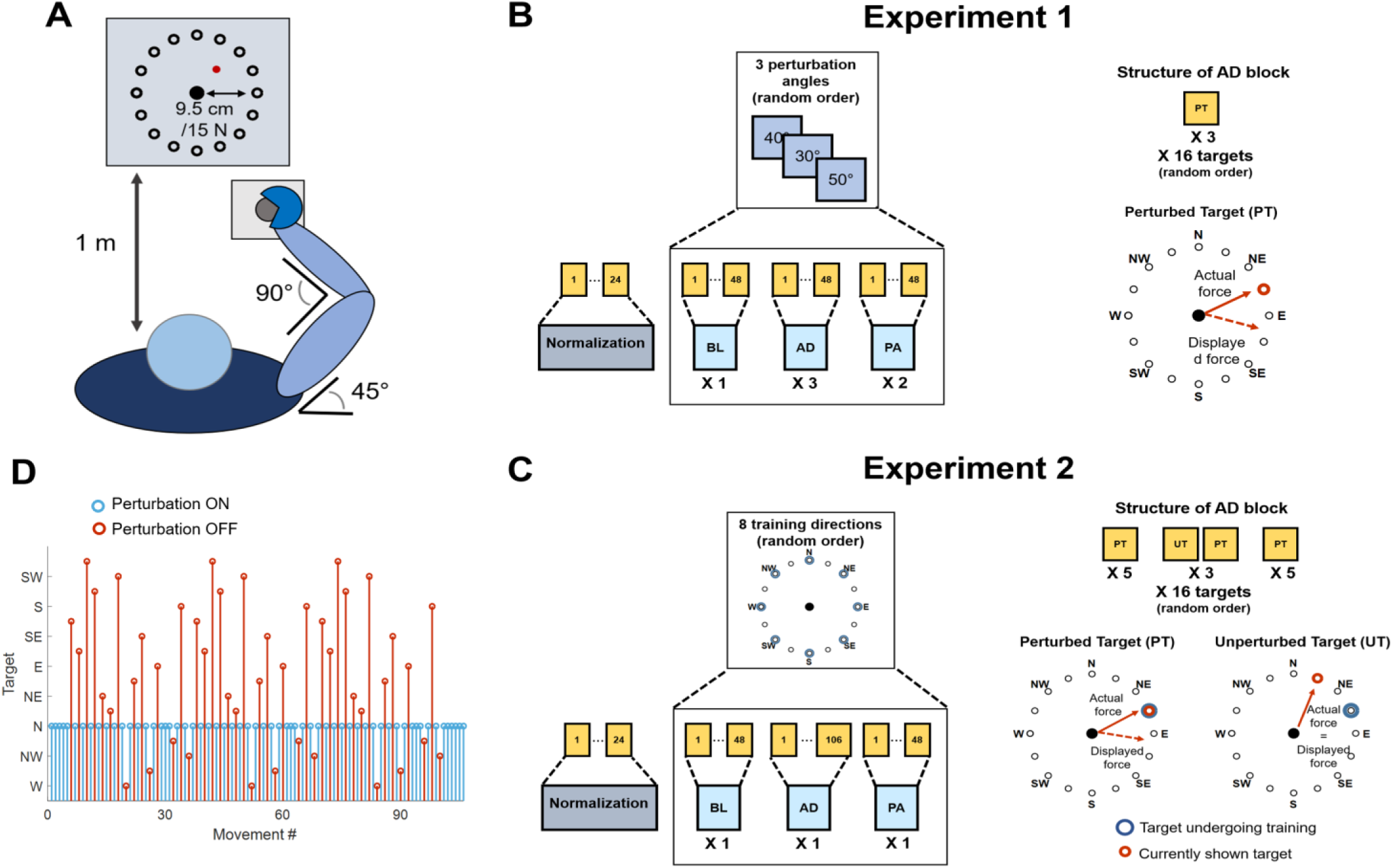
Experimental setup and procedures. **(A)** Graphical representation of the task that was employed in both experiments. Subjects kept their position consistent across all trials. The forearm was strapped to a support surface (not shown in the picture) and the hand was strapped to the manipulandum to avoid the use of the hand muscles during the task. Subjects were presented a virtual scene on a screen in front of them (1 m distance). The virtual scene consisted of a cursor, controlled in position by the force exerted on the manipulandum, and 16 targets, spaced 22.5° apart. **(B)** Protocol for Experiment 1. Subjects experienced a total of 19 blocks consisting of a normalization block (24 movements) and 3 macro-blocks of 6 block each, divided in baseline (BL, 1 block, unperturbed), adaptation (AD, 3 blocks, perturbed) and post-adaptation (PA, 2 blocks, unperturbed). Each block consisted of 48 movements. Each macro-block was characterized by a different clockwise (CW) rotation angle applied during the AD blocks (30°, 40° or 50°). In the AD blocks subjects experienced 3 repetitions of each target in a random order. The rotation was applied to all targets. **(C)** Protocol for Experiment 2. Subjects experienced a total of 25 blocks consisting of a normalization block (28 movements) and 8 macro-blocks of 3 block each, divided in baseline (BL, 1 block, unperturbed), adaptation (AD, 1 block, perturbed) and post-adaptation (PA, 1 block, unperturbed). The BL and PA block consisted of 48 movement. The AD block consisted of 106 movements. During the AD block the perturbation was applied to one target only (perturbed target, PT), while the mapping between force and cursor position was unperturbed for the other targets (unperturbed targets, UT). Each macro-block was characterized by a different perturbed target (among 8 different random targets, spaced 45° apart). Subjects first experienced the PT 5 times, then alternated between the PT and all the UTs in a random order for 3 times (for a total of 96 movements) and then concluded the block with 5 consecutive repetitions of the PT. **(D)** Graphical representation of the target order experienced during the AD phase of Experiment 2. In blue is presented the perturbed target (in this case N), in red the unperturbed ones.

**Experiment 1** consisted of 19 blocks (**Figure 1B**). The first block consisted of a normalization block that was used to determine the average EMG activity relative to 8 reaching directions covering the whole workspace at angular intervals of 45°. During the normalization block subjects were asked to reach for each one of the eight targets (presented in a random order) and hold the cursor on the target for 5 seconds. Subjects repeated the reach-and-hold task three times for each target, for a total of 24 movements. The following 18 blocks were divided in 3 macro-blocks each constituted by 6 blocks. In each macro-block, subjects experienced 1 baseline block (BL), where they were asked to reach for all the 16 targets three times (48 total movements) without the visual perturbation. Subjects then experienced 3 adaptation blocks (AD1, AD2 and AD3) where they reached for all the 16 targets three times (48 total movements) while the visual perturbation was applied to the whole workspace. Finally, subjects experienced 2 post-adaptation blocks (PA1 and PA2), where they were asked to reach for all the 16 targets three times (48 total movements) without the visual perturbation. Each macro-block was characterized by a different visual perturbation angle during the AD blocks, equal to 30°, 40° or 50°, in a random order. All 3 AD blocks of a macro-block were characterized by the same visual perturbation angle.

**Experiment 2** consisted of 25 blocks (**Figure 1C**). The first block of Experiment 2 consisted of a normalization block, identical to the one experienced in Experiment 1. The following 24 blocks were divided in 8 macro-blocks each constituted by 3 blocks. During each macro-block subjects experienced a baseline block BL identical to the one experienced during Experiment 1 (48 unperturbed movements, 3 per target in a random order). Then subjects experience an adaptation block AD, consisting of 106 reaching movements, where a 45° visual perturbation was applied only to one target, while the virtual scene was unperturbed for the other 15 targets (**Figure 1C and D**). Subjects were first asked to reach for the perturbed target 5 times, then they were asked to reach for all the 16 targets (including the perturbed one) three times, each repetition interspersed by a single repetition of the perturbed target. Thus, each reaching movement to one of the 16 targets, presented in a random order, was followed by a movement to the perturbed target. Subjects in this phase alternated perturbed and unperturbed movements except for when the perturbed target was interspersed with itself, where they experienced 3 consecutive perturbed targets. Subjects concluded the block by experiencing the perturbed target 5 consecutive times. In total, during the AD block, subjects performed 45 unperturbed and 61 perturbed movements (an example of the order of perturbed and unperturbed targets in the AD block is presented in **Figure 1D**). The design of this block allowed for evaluating how adapting for a perturbation acting on one single target affected also the reaching to the unperturbed targets. At the same time, this experimental design counteracted the forgetting effect that reaching for unperturbed targets has on the adaptation process. Each of the 8 macro-blocks was characterized by a different perturbed target during the AD block. After the AD block, subjects experienced a single PA block, identical to the ones experienced during Experiment 1. The perturbation was applied to 8 targets covering the whole workspace at angular intervals of 45° (**Figure 1C**). The order of the perturbed target, and thus of the macro-blocks, was randomized.

### Analysis of reaching movements

Data from the load cell were filtered using a low-pass filter (Butterworth, 3^rd^ order) with cut-off frequency set at 10Hz. Changes in the force trajectories during the different phases of both the experiments were characterized using the initial angular error (IAE) metric. The IAE was calculated (**Figure 2**) as the angle between the straight line connecting the center of the workspace with the intended target and the straight line connecting the center of the workspace with the actual position of the cursor at 2.6 cm from the center (equivalent to 4 N of force exerted) during each movement. This distance was selected based on the data-driven observation (**Figure 3A, B** and **C** and **Figure 4A**) that subjects started compensating for the initial angular errors only after about half of the movement trajectory (equivalent to 7.5 N), thus the metric allows to capture a point in time where the subject is “committed” to the movement but has not yet started compensating for the initial shooting error. In the analysis of Experiment 2, we analyzed the IAE metric as a function of the distance between the target analyzed and the perturbed target. In this analysis, we pooled together the data relative to the AD phase of each macro-block and we calculated the average (across macro-blocks and subjects) IAE for each target as a function of their angular distance from the perturbed target. Moreover, we analyzed the behavior of the IAE metric both for the repetitions of the perturbed target only and for the repetitions of its 4 (2 clockwise, 2 counterclockwise) closest targets.

**Figure 2.**
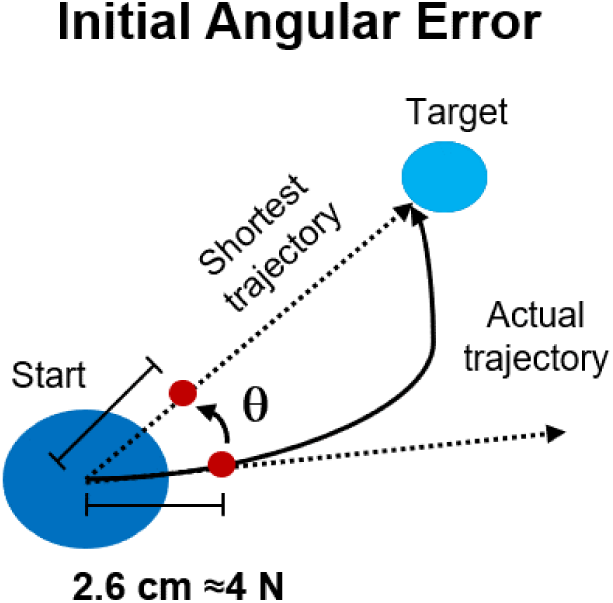
Performance metrics for reaching in both experiments. The initial angular error was calculated, for each movement repetition, as the angle between the optimal, shortest, straight trajectory and the actual trajectory at 2.6 cm from the center of the workspace.

**Figure 3.**
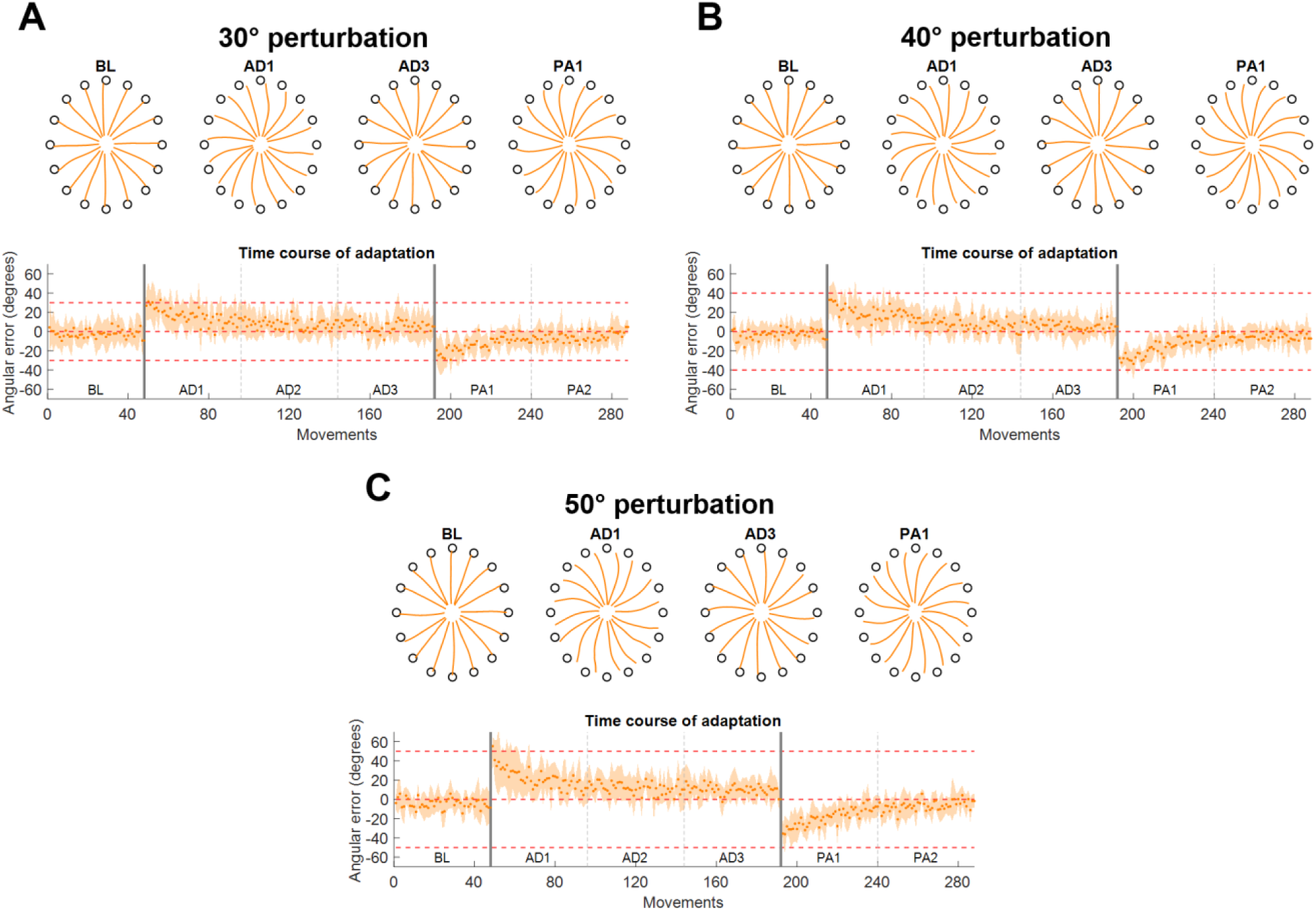
Force trajectories and initial angular error (IAE) results for Experiment 1. Each panel presents the results for a different perturbation angle (A for 30°, B for 40° and C for 50°). Each panel presents, on the top plot, the average (across subjects and repetitions) force trajectories for the last 5 movements of BL, the first 5 movements of the first block of AD (AD1), the last 5 movements of the last block of AD (AD3) and the first 5 movements of the first block of PA (PA1). The bottom plot presents the average (across subjects) values of IAE for each movement across all blocks. The two vertical grey lines represent the onset and offset of the visual rotation. Horizontal red dotted lines represent the angle of the perturbation.

**Figure 4.**
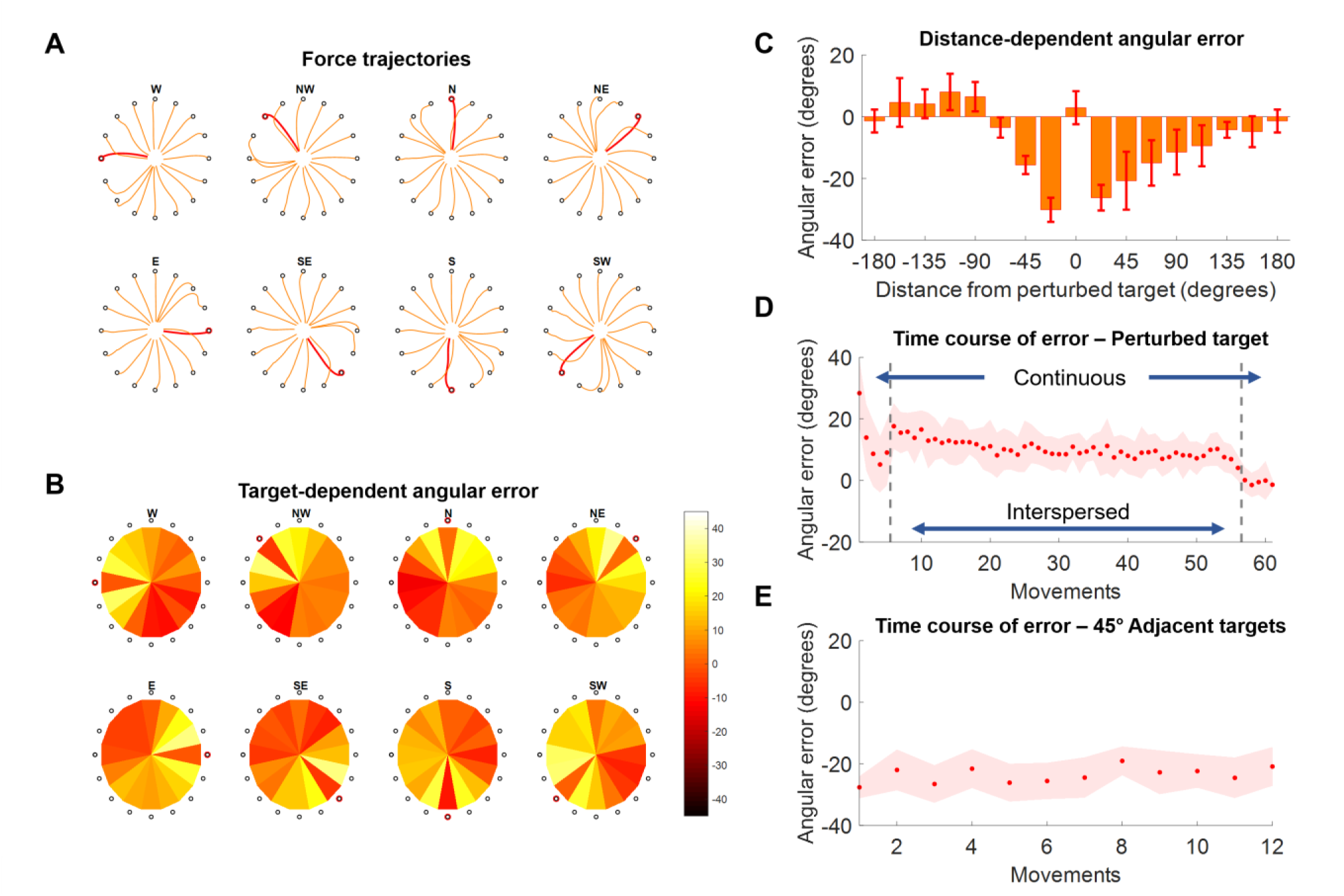
Force trajectories and initial angular error (IAE) results for Experiment 2. **(A)** Force trajectories for the last 5 movements of each target during AD, for each perturbed target. Trajectories for the perturbed target are in red. **(B)** Average values of IAE for the last 5 movements of each target during AD, for each perturbed target (indicated by a red circle). Each pie chart presents the average across all subjects. **(C)** Distribution of average (across subjects and targets) IAE values for the last 5 repetitions of each target grouped with respect to the distance between the target and the perturbed one (were 0 indicates the perturbed target itself). **(D)** Average (across subjects) IAE values for all the perturbed targets across all the repetitions of the AD block. During the first and last 5 repetitions the perturbed target is presented continuously, while in the middle section of the experiment (denoted by the two vertical grey dashed lines) the perturbed targets are presented interspersed with all the other targets. **(E)** Average (across subjects) IAE values of the 4 targets between −45° and 45° of the perturbed one, in order of occurrence (12 total occurrences).

### EMG signal recording and processing

EMG signals were recorded, during both experiments, from the following 13 upper limb muscles: Brachiradialis (BRD), Biceps brachii short head (BSH), Biceps brachii long head (BLH), Triceps brachii lateral head (TLT), Triceps brachii long head (TLN), Deltoid Anterior (DANT), Medial (DMED) and Posterior (DPOST) heads, Pectoralis Major (PM), Inferior head of the Trapezius (TRAP), Teres Major (TMAJ) and Latissimus Dorsi (LD). EMG signals were recorded through a Delsys Trigno system (Delsys, US), sampled at 2000 Hz and synchronized with the load cell. EMG signals were first filtered in the 20Hz-400Hz band by using a 3rd order digital Butterworth filter. The envelopes were then obtained by rectifying the signals and applying a low pass filter (3rd order Butterworth) with a cut-off frequency of 10Hz. Before muscle synergies extraction, all the envelopes were amplitude normalized. The normalization was done with respect to the subject- and session-specific reference values calculated from the normalization block. During the normalization block, subjects reached three times to 8 targets spaced at 45°. The target associated with the maximal activation of each muscle was identified. The reference normalization value for each muscle was calculated as the average peak envelope value across the three repetitions of the target maximizing the muscle’s activity.

### Semi-fixed synergies model and synergy extraction

In the muscle synergies model, a matrix *M* containing *s* samples of the envelopes obtained from the EMGs recorded from *m* muscles is decomposed, using the non-negative matrix factorization (NMF) algorithm (Lee and Seung, 2001), as the combination of *n* muscle synergies *M* ≈ *W* · *H*, where *W* represent a matrix of *m* · *n* synergy weights and *H* represents a matrix of *n* · *s* synergy activation patterns.

We and others have shown (Gentner et al., 2013; De Marchis et al., 2018; Zych et al., 2019) that adaptations to perturbations in several different tasks are well represented by the changes in the activation patterns *H* of fixed sets of muscle weights *W* extracted by applying the NMF algorithm to sets of EMG signals recorded during unperturbed versions of the tasks under analysis. This analysis is usually performed by altering the NMF algorithm by fixing the values of *W* while allowing the update rule of the NMF algorithm to modify only the values of *H*. The validity of the fixed-synergies model is often evaluated by showing that the EMG reconstructed using the fixed set of *W* and the new *H* can capture the variance of the data up to an arbitrary satisfactory level of a performance metric (e.g. 90% of the variance accounted for).

There are some conceptual and technical limitations to the fixed-synergies approach. In first instance, this model requires that the muscle synergies are fully represented, at the neurophysiological levels, by the matrix *W*, which hard codes the relative activations of the different muscles relative to each synergy module. Even if the neurophysiological muscle synergies were consistent with this spatially fixed synergistic model (rather than, e.g., a dynamic synergy model such as the ones described in (d’Avella et al., 2003) and (Delis et al., 2014)), it is unlikely that the relative activation of the different muscles would be hard-fixed, but rather “stabilized” by the neurophysiological substrates encoding the synergies. We found, in fact, that single muscular activations can be altered, within the synergies, depending on task demands (Zych et al., 2019).

Moreover, a technical limitation of the standard fixed-synergies approach lies in the fact that EMG recordings can undergo changes in conditions during a recording session (e.g. sweat during long tasks can alter the signal-to-noise ratio of a channel) and between recording sessions, thus by fixing the relative weights between the muscles we may lose variance in the reconstructed data caused by exogenous, rather than endogenous, changes in the EMGs. For these reasons we here introduce the semi-fixed synergies model. In this model, the synergy weights *W*^*BL*^ extracted during an unperturbed baseline task are used to determine the range over which the single muscle contributions to the synergy weights extracted during adaptation can vary. Specifically, given:

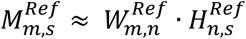

With 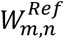 and 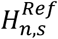 respectively the synergy weights and activation patterns extracted by applying the NMF algorithm on a reference (unperturbed) dataset, with the matrices 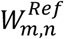 and 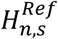 appropriately scaled so that 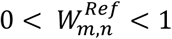, and given a weight tolerance *δ*, indicating the variability allowed around the values of 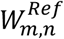 during the extraction of the muscle synergies for the adaptation/post-adaptation conditions, the semi-fixed synergies model bounds the results of the standard multiplicative update rule of the NMF on the weights so that:

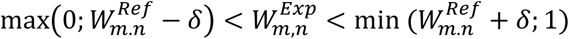

Thus, in the semi-fixed synergies model, the weights of the muscle synergies extracted during the different experimental phases are not fixed but bounded around the values of the weights extracted during the reference part of the dataset. The values of 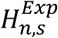 are left completely free to change, as in the fixed-synergies model. In the semi-fixed model most of the variability of the data between a baseline and an adaptation/post-adaptation condition is described by changes in the synergy activation patterns, while a smaller part of such variability is ascribed to changes in the weights.

In all our subsequent analyses, the value of δ was fixed to 0.1, meaning that the weights of the individual muscles in a synergy were allowed a 10% variability in the positive and negative directions with respect to their values in the reference synergy weights. This value was chosen to capture the variability of the muscular weights in a context (isometric movements in a fixed posture) where small variability is expected. In the analysis of Experiment 1, the reference *W*^*Ref*^ was calculated from the data pooled from the BL blocks relative to the 3 macro-blocks. The EMG envelopes calculated from each BL block were concatenated in temporal order and then smoothed using a 4-points average filter, in order to avoid hard transitions between the data of the different BL blocks. Similarly, in the analysis of Experiment 2 the reference *W*^*Ref*^ was calculated from the data pooled from all the 8 BL blocks relative to the 8 different macro-blocks, following the same procedure as for Experiment 1.

After the extraction of the reference synergies, the semi-fixed *W* and *H* were extracted from all the experimental blocks of both experiments (including the single BL ones) using the procedure for semi-fixed synergies extraction previously described. In all our analyses, the number of muscle synergies extracted was fixed to 4. This number of synergies was found by us and others (Berger et al., 2013; Gentner et al., 2013; De Marchis et al., 2018) to well represent the variability of the upper limb muscular activations during planar isomeric reaching movements. Moreover, the 4 synergies have been shown to have distinct activation sub-spaces (as determined by the RMS of the activation of each synergy relative to each target, see later) that heterogeneously cover the whole planar workspace, with each synergy spanning approximately 90° (De Marchis et al., 2018).

We evaluated the quality of the envelope reconstruction obtained in each block using the semi-fixed synergy model by calculating the R^2^ between the original envelopes and the envelopes obtained by multiplying *W*^*Exp*^ and *H*^*Exp*^. To assess for statistically significant differences in R^2^ across the different blocks we employed ANOVA for comparing the average (across macro-blocks) R^2^ obtained in each block, for both experiments. Finally, in order to justify subsequent group analyses on the synergy activations, we evaluated the similarity between the *W*^*Ref*^ extracted from each subject using the normalized dot product. In order to do so, we calculated, for each subject, the similarity between the *W*^*Ref*^ matrix of the subject and the *W*^*Ref*^ matrices of all the other subjects and then averaged it, so to obtain a subject-specific similarity measure.

### Synergy and muscle rotation analysis

Previous works have shown that adaptations to visuomotor rotations during planar isometric movements are well described by rotations of the sub-spaces where the different synergies and muscles are active in the overall workspace (Gentner et al., 2013; De Marchis et al., 2018). Here we employed the same analysis in both experiments in order to characterize how adapting to different perturbation angles (Experiment 1) and in different sub-spaces (Experiment 2) modifies the activation patterns of the muscle synergies. In order to do so we first estimated the workspace covered by each of the synergies in each experimental block.

This was done by: i) segmenting the *H* matrix calculated for each block by extracting the sub-portion of *H* relative to the center-out phase of each reaching movement, from the instant when the target appeared on screen to the instant when the target was reached; ii) calculating the RMS of the *H* for each reaching movement; iii) averaging the values of RMS across the different repetitions of each target in a block. For all blocks (BL, AD and PA of each macro-block) in Experiment 1 and for the BL and PA blocks in Experiment 2 the average was calculated across all three repetitions of each target. For the AD block of Experiment 2, the RMS values relative to the unperturbed targets were also averaged across all three target repetitions in the block, while those relative to the perturbed target (which the subjects experienced 61 times in the training block) were averaged across the last 3 interspersed repetitions that they experienced in the block before the final 5 continuous ones. This choice was suggested by the results obtained while analyzing the biomechanical characteristics of adaptation in Experiment 2 (**Figure 4D**), that showed that subjects had reached adaptation during the final part of the interspersed trials, while still showing the influence of the presence of the non-perturbed trials.

We then calculated the preferred angle spanned by the activation pattern of each single synergy in the workspace (d’Avella et al., 2006). Preferred angles were calculated from the parameters of a cosine fit between the average RMS of each synergy activation and the corresponding target position. RMS values were fitted using a linear regression in the form: *RMS*(*θ*) = *β*_0_ + *β*_1_ cos(*θ*) + *β*_2_ sin(*θ*). The preferred angle of the fit was then calculated from the fitting parameters as *ϑ* = *tan*^−1^(*β*_2_/*β*_1_). Only preferred angles calculated from significant (p < 0.05) fittings were used in subsequent analyses. In both experiments we evaluated the difference in preferred angles between the BL blocks and the different AD and PA blocks. We refer to these differences as the rotations in preferred angles, or tunings, due to the adaptation process.

In Experiment 1, we analyzed the rotation of each synergy for each subject during all the AD and PA blocks of each macro-block. Moreover, we also evaluated the rotation of the average (across subjects) *RMS(θ)* of each synergy at AD3 for all three perturbation angles.

In Experiment 2, in each macro-block, we analyzed the rotation of each synergy of each subject for each perturbed target during AD. We grouped the rotations relative to the adaptations to the different perturbed targets depending on the angular distance between the perturbed target and the preferred angle of each synergy. We did this both across all perturbed targets and synergies and for each perturbed target singularly by ranking the synergies from the closest to the furthest to the perturbed target in terms of absolute angular distance with the synergy preferred angle.

Finally, as a validation of our approach, we calculated the preferred angles also for each of the 13 muscles and then calculated the rotations that these preferred angles incurred between BL and AD3 in Experiment 1 and between BL and AD for Experiment 2, using the same procedures we employed for the synergies activation patterns. We then assessed if the rotation of the single muscles correlated with the rotation of the synergies to which they contribute. A muscle was considered as contributing to a synergy if its weight in the synergy was above 0.25 (De Marchis et al., 2015) where, in our model, the maximum value that a muscle can have in a synergy is 1. We evaluated the correlation using Pearson’s coefficient, applied to the data pooled across subjects, synergies and experiments.

## Results

### Force Trajectories

The results on the analysis of the force trajectories and the IAE metric for Experiment 1 followed closely the results obtained in literature in similar experiments (Krakauer et al., 1999; Krakauer et al., 2000; Wigmore et al., 2002; Gentner et al., 2013). Across the three perturbation angles, we found that subjects, on average, presented increasing values of IAE with increasing perturbation angles in the first movement of the first AD block (26.9 ± 15.3°, 33.0 ± 14.0° and 55.4 ± 9.7° for the 30°, 40° and 50° perturbations respectively) and they were subsequently able to adapt and come back to a smaller IAE (<7° on average in the last 5 movements of each AD3 block for all three perturbations) through the repetitions of the different movements in the three AD blocks (**Figure 3A, 3B** and **3C**). The adaptation exhibited an exponential behavior.

In Experiment 2 we found that subjects were able to adapt their force trajectories to perturbations applied to a single target (**Figure 4A**). Subjects were able to minimize the IAE metric for the trained target, and this was mirrored by an IAE opposite to that induced by the perturbation in the adjacent, unperturbed, targets (**Figure 4B**). We found that targets positioned both clockwise and counterclockwise with respect to the perturbed target were affected by the adaptation and presented rotations opposite in direction with respect to the angle of the visual perturbation (**Figure 4C**). Targets positioned clockwise with respect to the perturbed target presented substantial counter-rotations up to about 120° of angular distance to the perturbed target, while the same effect was present counterclockwise only up to about 70° of angular distance (**Figure 4C**).

At the temporal level, the perturbed targets first exhibited a decrease in IAE metric during the 5 continuous movements at the beginning of the AD trial (**Figure 4D**). The average values of IAE increased as subjects began to experience the unperturbed targets interspersed with the perturbed one. Nevertheless, they were able to compensate for the presence of the unperturbed targets and reached an average value of IAE <10° by the end of the interspersed phase. They were finally able to reach an IAE value close to 0° during the last 5 continuous perturbed movements. On the other hand, the 4 45°-adjacent targets (2 clockwise and 2 counterclockwise) presented a constant average IAE value (about 25° of counterclockwise rotation) across their 12 repetitions (3 per target), indicating that the effect of the adaptation for the perturbed target over the unperturbed ones was maintained constant over the AD block (**Figure 4E**).

### Synergy extraction and validation of the semi-fixed synergy model

Consistently with what we previously showed (De Marchis et al., 2018), we found that 4 synergies can well represent the activity of all the muscles during both experiments. The 4 synergies were distinctly distributed in the different quadrants of the workspace and presented consistent preferred angles across the different subjects. In the following the preferred angles will be indicated using the W target (in a compass rotation) as 0° and increasing clockwise and the workspace will be referenced to by using the terms far and close for the upper and lower parts and lateral and medial for the left and right parts of the workspace, using the right arm as reference (**Figure 5A** and **5D**).

**Figure 5.**
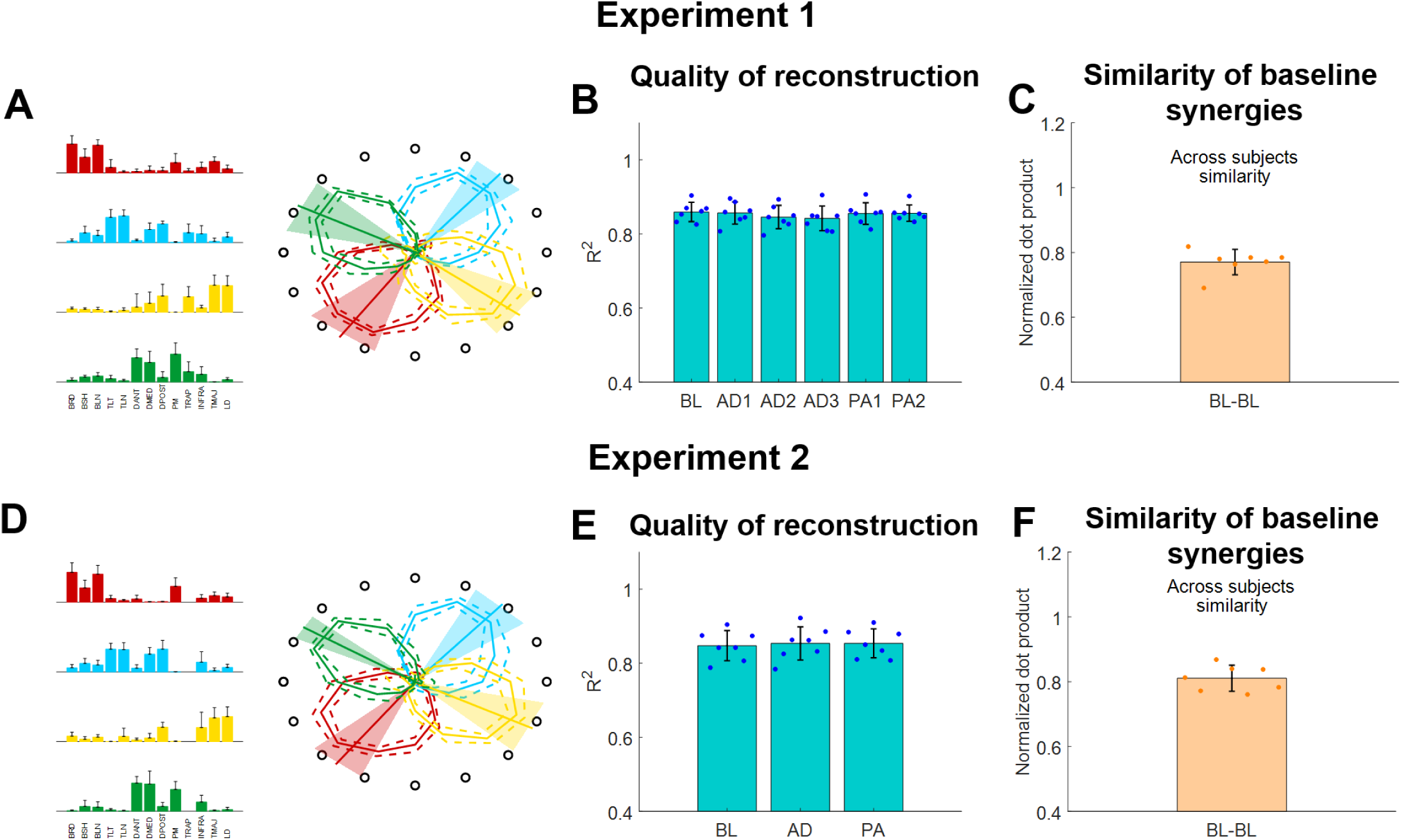
Muscle synergies extracted using the semi-fixed algorithm for both experiments. (**A** and **D**) Baseline synergy weights (average and standard deviations across subjects) and preferred angles across the workspace (bold line represents the average across subjects, shaded areas represent the standard deviation). (**B** and **E**) R^2^ of reconstruction for the synergies extracted from each block using the semi-fixed algorithm. Blue dots indicate the values of each individual subjects (averaged across macro-blocks), bars and whiskers indicate the average across subjects and the standard deviation. (**C** and **F**) Similarity of baseline synergies across subjects. Each dot represents the average similarity between one subject and all the other subjects. Bar and whiskers indicate the average across subjects and the standard deviation.

The synergies will be referenced-to using the color-coding of **Figure 5**. The red synergy was characterized by the activation of the elbow flexors and was active in the close-medial quadrant of the workspace. This synergy presented a preferred angle of 305.1 ± 17.3° for Experiment 1 and 307.1 ± 12.9°for Experiment 2. The green one synergy was characterized by the activation of the deltoids (medial and anterior), pectoralis and trapezius and was mostly active in the far-medial quadrant of the workspace. This synergy presented a preferred angle of 130.4 ± 12.4° for Experiment 1 and 131.6 ± 14.1° for Experiment 2. The azure synergy was characterized by the activation of the triceps, deltoid posterior and infraspinatus and was mostly active in the far-lateral quadrant of the workspace. This synergy presented a preferred angle of 217.3 ± 14.4° for Experiment 1 and 206.8 ± 15.1° for Experiment 2. The yellow synergy was characterized by the activation of the latissimus dorsi and teres major and was mostly active in the close-lateral quadrant of the workspace. This synergy presented a preferred angle of 26.9 ± 15.0° for Experiment 1 and 15.8 ± 7.1° for Experiment 2 (**Figure 5A** and **5D**).

The 4 synergies were able to well describe the variability of the data for the reference datasets (obtained, in both experiments, by pooling together the data of the BL blocks). We observed an average (across subjects) R^2^ of 0.86 ± 0.04 for the reference synergies extracted during Experiment 1 and an average R^2^ of 0.84 ± 0.05 for the reference synergies extracted during Experiment 2. When analyzing the average (across subjects and macro-blocks) R^2^ for the different experimental blocks as reconstructed using the semi-fixed synergies algorithm from the reference synergies, we found that the R^2^ values were above 0.8 for all blocks in Experiment 1 (**Figure 5B**). Moreover, we did not observe statistically significant differences in the R^2^ values among the different blocks (p = 0.98, ANOVA 1-way). The same results were observed also for Experiment 2 (**Figure 5E**), were the data reconstructed using the synergies extracted using the semi-fixed approach maintained an average (across subjects and macro-blocks) R^2^> 0.8, with no statistically significant differences across the different blocks (p =0.99, ANOVA 1-way).

Finally, we analyzed the across-subjects similarity between the reference baseline synergies calculated for each subject. We found an average similarity of 0.77 ± 0.04 for Experiment 1 and of 0.81 ± 0.04 for Experiment 2, indicating that subjects have similar synergies among them in both experiments.

### Synergies Rotations

In this analysis we evaluated how the workspace spanned by the activation patterns of each synergy changed during the different adaptation exercises. In Experiment 1 we found that, for all three perturbation angles, the synergies rotate almost solitarily (**Figure 6A**) by angles close to the one of the visual perturbations (**Figure 6B, 6C** and **6D**). These results are in line with what presented in (Gentner et al., 2013), where the author showed that a 45° visual rotation induces a rotation of the activation pattern of the synergies close to 45°.

**Figure 6.**
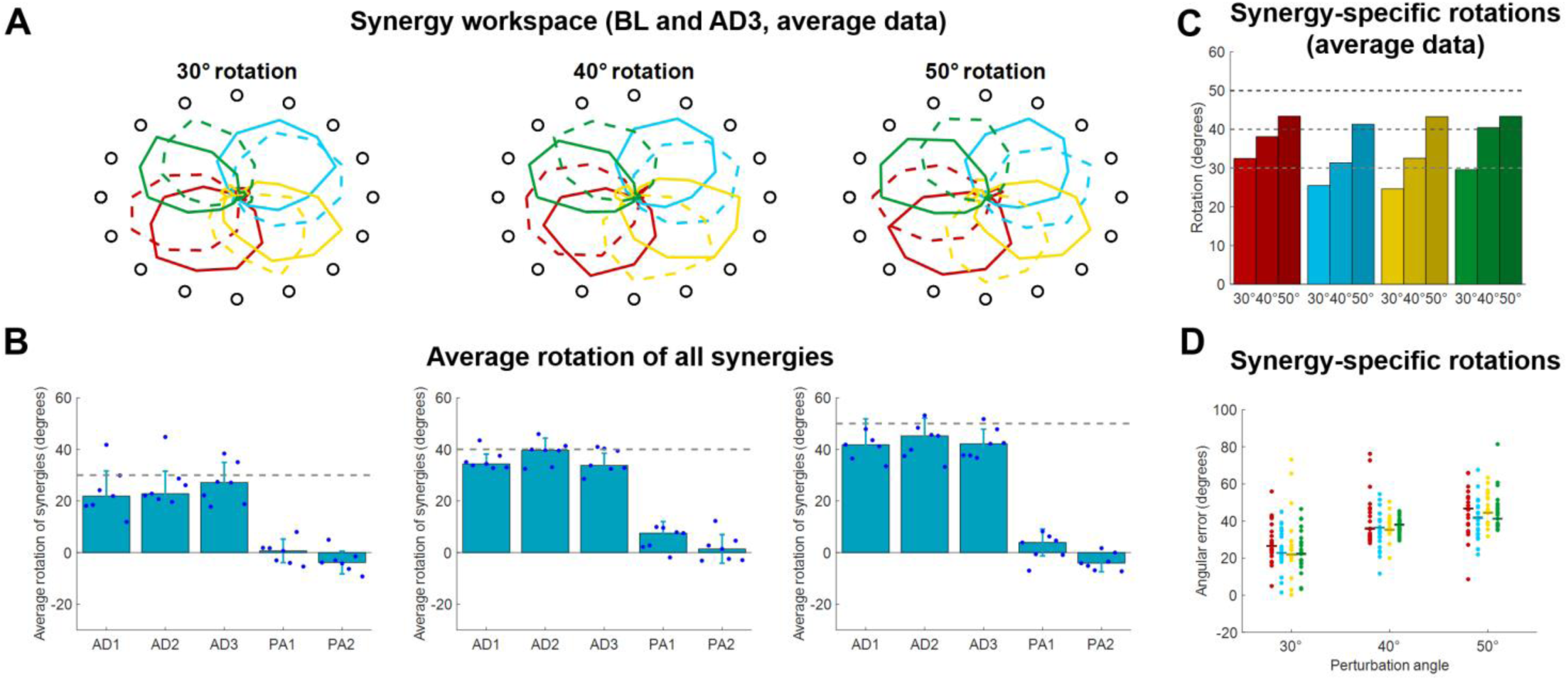
Synergies rotations for Experiment 1. **(A)** Average (across subjects) *RMS(θ)* of synergies activations for each target for BL (solid lines) and AD3 (dashed lines) for all three perturbation angles. **(B)** Average synergies rotation, with respect to their preferred angles at BL, for each block in each macro-block. Individual dots represent the data for each subject, as average rotations of all the 4 synergies. Bars and whiskers represent the average and standard deviation across subjects. The dashed grey lines represent the angle of the visual rotation. **(C)** Rotations at AD3 for each synergy in each macro-block, calculated from the average (across subjects) intensity of synergy activation (as in **A**). (**D**) Rotations at AD3 for each synergy in each macro-block calculated for each single subject (dots). The horizontal lines indicate the median rotation across subjects.

We analyzed the average (across synergies) rotation of the synergy workspace for each subject in each block (**Figure 6B**). Here we observed that subjects, across the three perturbations, appear to increase their average synergy rotation after the first block and achieve maximal rotation in the 3^rd^ (30° perturbation) or 2^nd^ (40° and 50° perturbations) block of adaptation. Subjects do not appear to show an after-effect in the synergies, but rather a small residual rotation. This result is expected and was previously observed in another adaptation study (Zych et al., 2019) and indicates that biomechanical after-effects such as the ones observed in **Figure 3** arise from the utilization of the adapted synergies in the unperturbed space.

For the rotations calculated from the average (across subjects) synergy *RMS(θ)* at AD3 (**Figure 6C**), we found rotations spanning from 24.6° (yellow synergy) to 32.5° (red synergy) for the 30° perturbation, 31.4° (azure synergy) to 40.4° (green synergy) for the 40° perturbation and 41.3° (azure synergy) to 43.4° (red synergy) for the 50° perturbation. We found similar results for the rotations calculated from the data of each single subject (**Figure 6D**), although subjects exhibited high variability among them for each combination synergy/perturbation-angle. We observed a range of median rotations spanning from 21.9° (yellow synergy) to 26.6° (red synergy) for the 30° perturbation, 35.5° (yellow synergy) to 36.8° (green synergy) for the 40° perturbation and 43.3° (green synergy) to 46.6° (red synergy) for the 50° perturbation.

In Experiment 2 we tried to characterize how the different synergies rotate when only a sub-space of the workspace is perturbed. An initial visual analysis of the average (across subjects) synergies *RMS(θ)* at BL and AD **(Figure 7**) sparked two initial observations: i) only the synergies involved in the reaching to the perturbed target are rotated in the adaptation process; ii) synergies whose preferred angle is close to the angle of the target being perturbed are not rotated. These two observations are equivalent to the observation that synergies are rotated only if engaged at the boundaries of their activation workspace.

**Figure 7.**
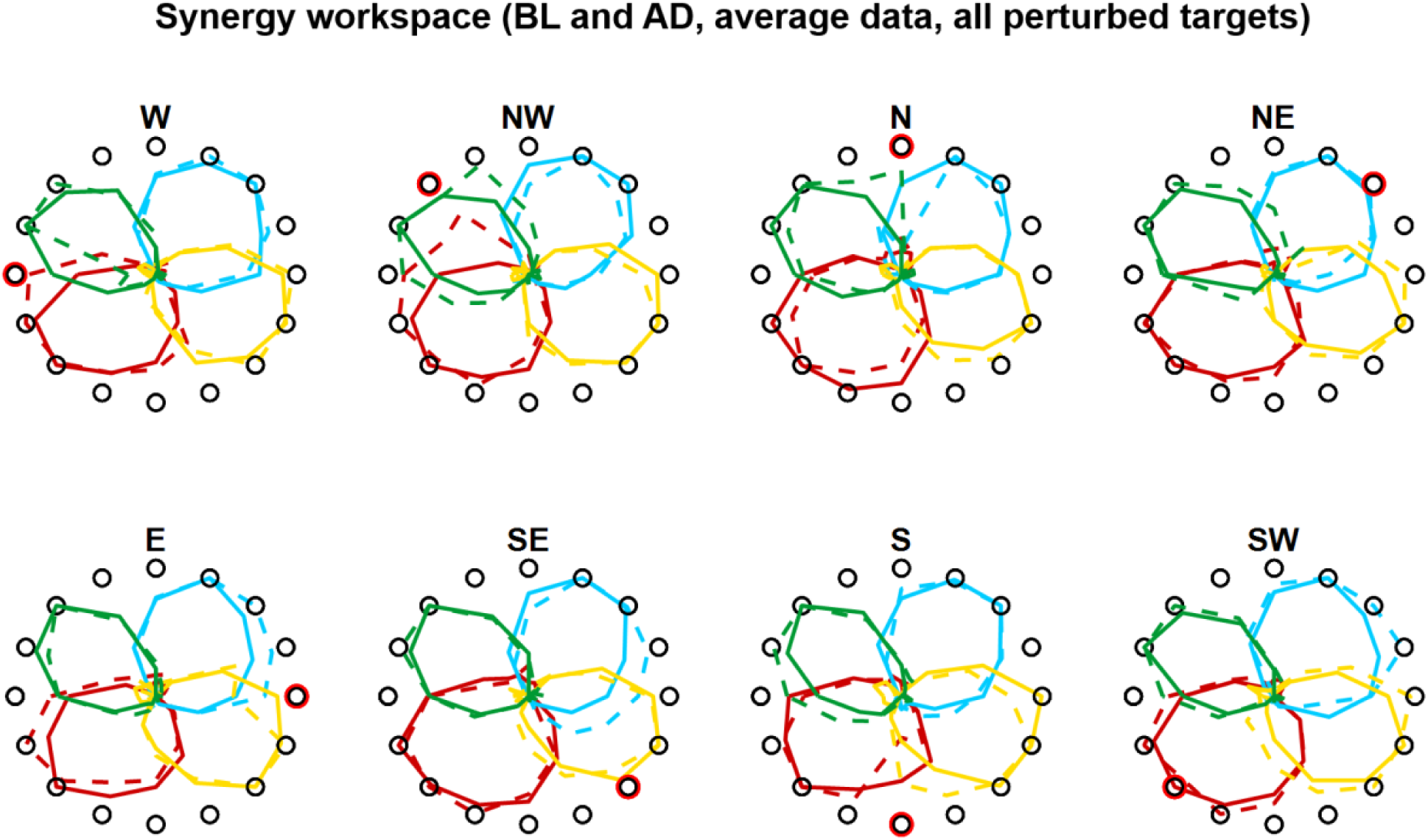
Synergies rotations for Experiment 1. **(A)** Average (across subjects) *RMS(θ)* of synergies activations for each target for BL (solid lines) and AD (dashed lines) for all perturbed targets. In the AD block, for the unperturbed targets the values are calculated from all three repetitions of each target, while the values for the perturbed targets are calculated from the last 3 repetitions during the interspersed phase of the block (see Fig. **1D** and **4D**)

The analyses of the synergy rotations of the single subjects confirm this observation. We observed that each synergy is maximally rotated during the adaptation to the perturbed target that is approximatively 90° clockwise with respect to the preferred angle of the synergy at baseline (**Figure 8A**). This observation is true for all 4 synergies, although they seem to exhibit different degrees of “sensitivity” to the adaptation process. In this regard, the azure synergy is only rotated for perturbed targets that are 45° to 120° clockwise with respect to the synergy preferred angle and the yellow synergy exhibits small values of rotation during almost all adaptation blocks. The analysis of the rotations for the 4 synergies pooled together further confirms the original observation (**Figure 8B**) and shows that the rotation of the synergies is close to 0° when the preferred angle of the synergy is very close (< 20°) to the perturbation angle. The rotation then increases in the clockwise direction reaching a maximum of about 20° at about 90° of distance between the perturbation angle and the synergy preferred angle and decreasing afterwards. In the counterclockwise direction, we observed an increase in rotation up to about a distance of 60° and inconsistent results afterwards.

**Figure 8.**
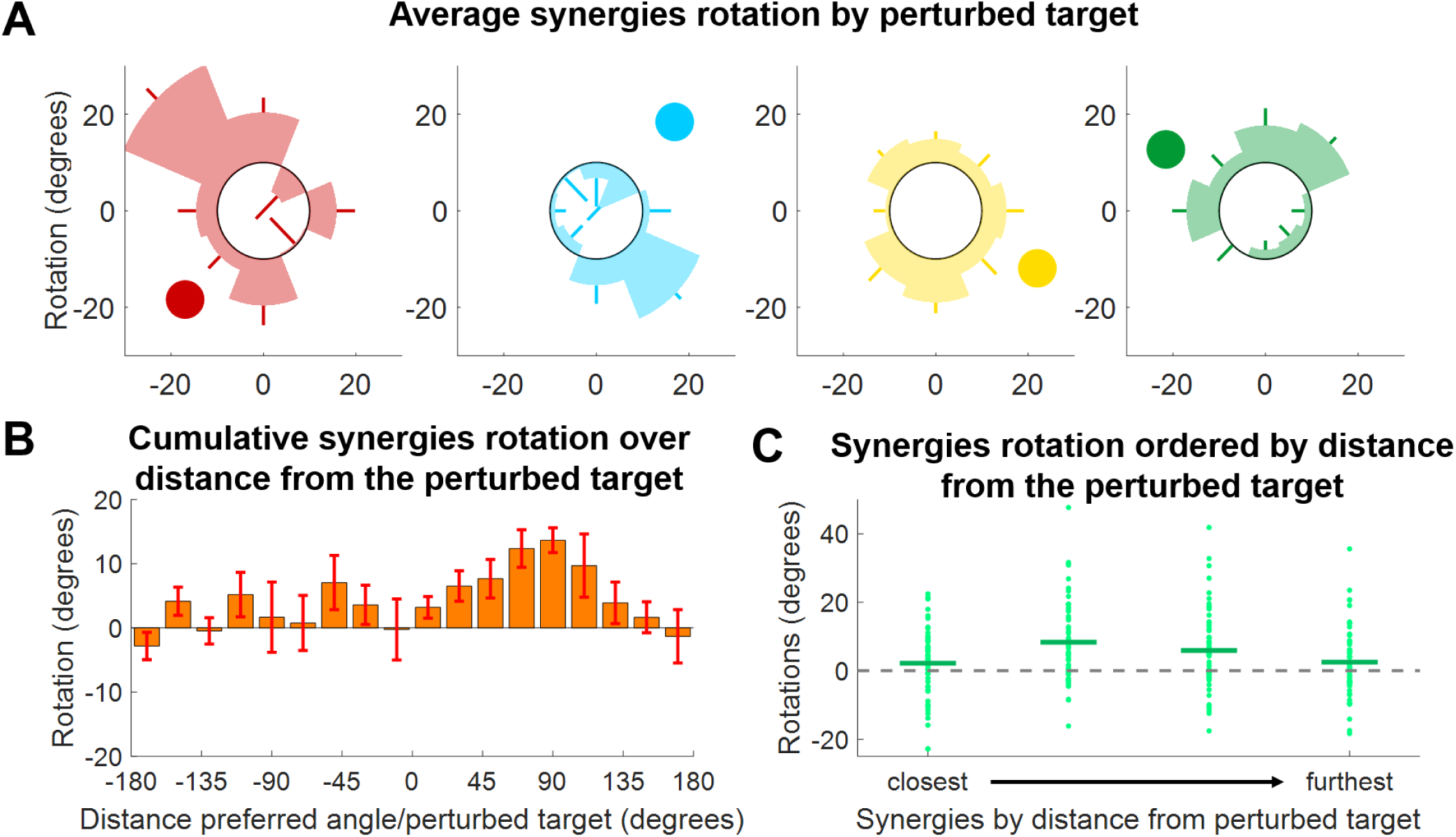
Synergies rotations for Experiment 2. **(A)** Average (across subjects) rotation for each synergy (color-coded) and for each perturbed target. Each segment of each polar plot represents a perturbed target. The darker circle represents the direction of the preferred angle for each synergy at BL. **(B)** Distribution of average (across subjects, targets and synergies) synergy rotation values as a function of the distance between the synergy preferred angle and the perturbed target. Bars represent averages, whiskers standard deviations. **(C)** Synergies rotations for each macro-block after ordering the synergies from the closest to the perturbed target to the furthest. Individual dots represent the rotation of each single synergy (56 total dots, 8 targets times 7 subjects). Horizontal lines represent the median across all the individual values.

As an additional analysis we ranked, for each perturbation angle, the synergies from closest to furthest in absolute angular distance to the perturbed target (**Figure 8C**). We observed, once again, that synergies closer to the perturbation angle exhibit the smallest rotation, while higher rotations are observed in the second and third closest synergies. In this analysis, it is also possible to notice the high variability exhibited by the rotations. This variability may be inherent to the phenomenon observed or derived from the methodology employed, where raw data are first factorized, then segmented and then fitted to a cosine fit, with each passage potentially introducing additional variability.

In order to validate our approach of analyzing adaptations in the synergies, rather than muscular, space, we analyzed how the single muscles rotate, on average, in both experiments. In Experiment 1, we found (**Figure 9A**) that the average rotation of the muscles increased with the perturbation angle, with average values across subjects equal to 24.6 ± 4.6, 29.6 ± 3.8 and 41.3 ± 3.5 for the 30°, 40° and 50° perturbations respectively. In Experiment 2, we once again analyzed the relationship between the muscle rotation and the distance between the baseline preferred angle (of the muscles in this case) and the angle of the perturbation, in a homologue of the analysis presented in **Figure 8B**. We found (**Figure 9B**) that muscular rotations held a behavior consistent with that observed in the synergies (**Figure 8B**) by which muscles with preferred angles close to the perturbed targets are not rotated during the adaptation, while rotations increase in the clockwise direction up to a maximum distance of about 90° to 110°. Counterclockwise we observed rotations only for angular distances between the preferred angle and the perturbation that are smaller than 60°, as in the synergies analysis. Finally, we compared the rotations of the single muscles with the rotation of the synergies to which those muscles contribute to. In this analysis (**Figure 9C**) we observed a moderate significant linear correlation between the rotation of the synergies and of the muscles, characterized by a value ρ = 0.57. We found that the angular coefficient of the line better fitting the data was equal to 0.59, indicating an overall underestimation of the rotation in the synergy-based analysis, that appears to depend mostly from an underestimation of negative rotations.

**Figure 9.**
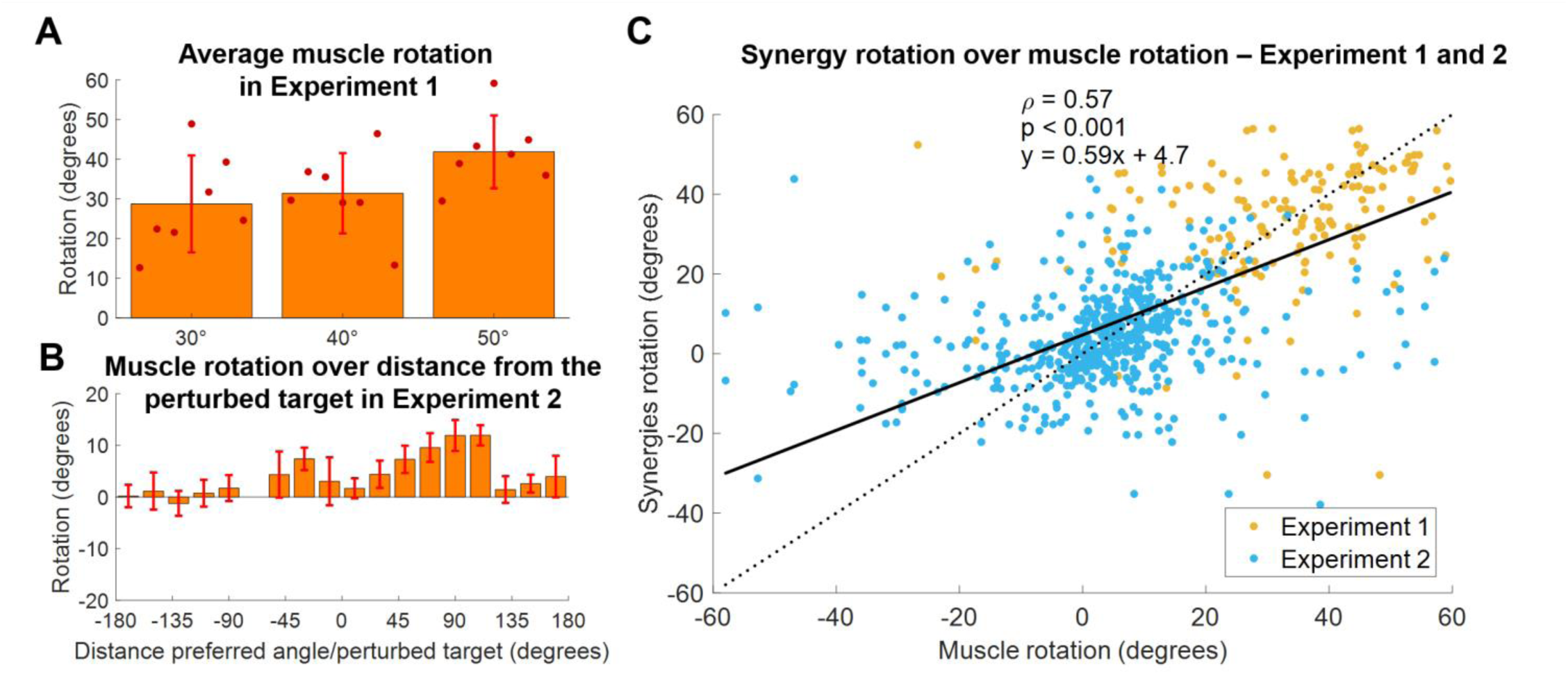
Comparison between synergies and muscle rotations. **(A)** Average (across muscles) rotation of the muscles at AD3 for all three macro-blocks of Experiment 1. Individual dots represent the average value for each subject in each experiment. Bars and whiskers represent the average and standard deviations across subjects. **(B)** Distribution of average (across subjects, targets and muscles) muscles rotations values as a function of the distance between the preferred angles of the muscles and the perturbed targets for Experiment 2. Bars represent average values, whiskers standard deviations. **(C)** Synergies rotations over the rotations of the muscles contributing to each synergy (data of both experiments pooled together). A muscle was considered to contribute to a synergy if its weight in the synergy was above > 0.4. The solid black line represents the linear fit between synergies and muscles rotations (values of the fit are presented in the plot, together with the ρ coefficient). The dotted line represents the fit relative to a perfect correspondence between muscles and synergies rotations.

## Discussion

In this study we sought to investigate how adaptations to visuomotor rotations are achieved in the neuromuscular space. We studied how muscular co-activations, modeled using muscle synergy analysis, are modified when different angular rotations are used to perturb the mapping between the force exerted and the visual feedback provided to the individuals during isometric contractions.

Specifically, we investigated how different rotations angles applied to the whole workspace and the same rotation applied to small sub-spaces modify the activations of the synergies. In our analysis we were particularly interested in identifying generalizable behaviors that could be potentially used to model the effect of a given visual perturbation on the neuromuscular control.

We found strong evidences supporting the observations that muscular activations and their synergistic homologues are tuned proportionally to the perturbation angle (**Figure 6** and **Figure 9A**) and only when engaged at the boundaries of their workspace (**Figure 7**), and with an angle proportional to the distance between the perturbed sub-space and the preferred direction of the muscle/synergy (**Figure 8** and **9B**). Our analysis shows that such behaviors are consistent whether analyzing muscular or synergies activations (**Figure 9B** and **9C**), further strengthening the argument that synergies analysis can simplify the description of adaptations to visuomotor rotations (Berger et al., 2013; Gentner et al., 2013; De Marchis et al., 2018).

In a previous work (De Marchis et al., 2018) we showed that adapting to perturbations affecting two sub-spaces of the whole workspace leads to different synergies rotations depending on the order in which the two perturbed sub-spaces are experienced. One of the aims of the work we present here was to investigate whether these differential neuromuscular paths to adaptation may depend on the relationship between the workspace covered by each single synergy and the spatial characteristics of the sub-space being trained.

Here we found evidences of such relationship that may help explain our previous results. In fact, we observed that the presence and extent of tuning in the synergies depend on the distance between the synergy preferred angle and the direction of the perturbed target.

Our results show that adapting for a 45° rotation applied to a sub-space does not lead to a precise 45° rotation of all the synergies, but leads to different rotations of the subset of synergies that are active in the sub-space, with the amount of rotation depending, for each synergy, on the spatial characteristics of the perturbed sub-space. In a scenario like the one we tested in our previous work (De Marchis et al., 2018), where two groups of subjects adapted for a 45° rotation applied to two sub-spaces experienced in opposite order, each group, after the first adaptation bout, achieved a different adapted neuromuscular state, as characterized by different tunings in the synergies. Therefore, each group had a different “starting” set of synergies preferred angles before the second adaptation bout and this could have led to the different “final” adapted states that we observed after adapting for the rotation applied on the second sub-space.

This interpretation of our previous results implies that the functional relationship that we identified between the preferred angles of the synergies and the workspace spanned by a visuomotor rotation could help to better understand some phenomena observed during visuomotor adaptations such as interference and transfer between adaptation processes. The first term refers to interference of prior adaptation to a subsequent adaptation process (Krakauer et al., 2005), while the second one refers to the generalization of a previously adapted behavior to a non-experienced scenario (Shadmehr, 2004). These two processes can be seen, at least functionally, as different aspects of the generalization of motor adaptations (Krakauer et al., 2006).

Visuomotor adaptation is a process involving the CNS at different levels starting from motor planning (Wong et al., 2015; Krakauer et al., 2019), and similarly, the processes driving generalization can also be traced at the motor planning level (Krakauer et al., 2006; Lerner et al., 2019), as exemplified also by studies that investigated the presence and extent of inter-limb generalization (Sainburg and Wang, 2002; Criscimagna-Hemminger et al., 2003; Wang and Sainburg, 2003). Nevertheless, several studies found that interference is task- and workspace-dependent (Bock et al., 2001; Woolley et al., 2007) and that generalization is constrained spatially to small sub-spaces of about 60°-90° degrees around the perturbed sub-space (Krakauer et al., 2000; Donchin et al., 2003; Brayanov et al., 2012). Thus, it appears that some aspects of the adaptation and generalization processes are dictated by biomechanical aspects, such as the workspace that the different actuators or actuating modules span in the movement space (de Rugy et al., 2009), up to the point where adaptations are only possible if they are compatible with the muscular activation space (Berger et al., 2013).

As an example, Wooley et al. (Woolley et al., 2007) showed that dual adaptation to opposing visuomotor rotations happens only when the workspaces associated with the two perturbations are different. When the opposing rotations are applied to the same workspace, the two adaptation processes interfere with each other. On the other hand, they showed dual adaptations to opposed rotations happening for targets that are 180 degrees apart. Interpreting their results in light of the ones that we show here suggests that the dual adaptation on disjointed workspaces can happen because different, non-overlapping synergies are involved in the process, while the dual adaptation on the same workspace is not attainable because it would require opposite rotations and counter-rotations of the same set of muscular modules.

An adaptation process constrained by neuromuscular coordination could perhaps also help explain the reference frame that is employed during visuomotor adaptation. It was generally assumed that visuomotor adaptation is performed in an extrinsic (world-based) reference frame (Krakauer et al., 2000), as also confirmed by studies on inter-limb generalization (Wang and Sainburg, 2004). Nevertheless, more recent studies suggested a mixed effect of adaptation in extrinsic and intrinsic (joint-based) coordinates (Brayanov et al., 2012; Carroll et al., 2014) and showed that adaptation to isometric tasks presents greater transfer in intrinsic coordinates (Rotella et al., 2015). The possibility that adaptation is biomechanically constrained by the muscle synergies (de Rugy et al., 2009) may explain this uncertainty of reference frame. In the muscle synergies space, intended in this case as the muscular coactivation maps that are semi-fixed in intrinsic coordinates (with variable individual muscular gains in each synergy that depend on task requirements (Zych et al., 2019)), an extrinsic adaptation at the motor planning level could generalize to an intrinsic reference frame by a magnitude proportional to the resultant of the synergies “tuning” (Gentner et al., 2013) in the intrinsic space (and vice-versa). This hypothesis, nevertheless, cannot be tested from our current dataset and requires a specifically designed experiment to confirm it.

Our results once again show the solidity of the synergy model in describing upper limb motor control and motor adaptations. This is relevant given the simplified biomechanical interpretational approach that the dimensionally smaller synergistic model allows with respect to the more redundant muscular space. Previous studies have shown that adaptation is obtained by tuning single muscles (Thoroughman and Shadmehr, 1999) and that this behavior is reflected (Gentner et al., 2013; De Marchis et al., 2018) in a spatially-fixed synergy model. It is not the aim of this paper to investigate whether the synergistic model, and in particular the static spatially fixed synergy model (as compared with other, more complex models (Delis et al., 2014)) well represents the neurophysiological structures that demultiplexes the cortical motor signals in the spinal cord. Our aim is rather that of understanding whether this relatively simple model can be used to describe visuomotor adaptations in a functional way, with potential applications aiming at the purposeful use of adaptations for obtaining desired kinematics and neuromuscular outputs, such as in the Error Augmentation scenario (Sharp et al., 2011; Abdollahi et al., 2014). However, such applications should consider also how the functional relationship herein identified at the neuromuscular level contribute to implicit and explicit processes of adaptation and learning (Taylor et al., 2014), given their differential effect on long term retention of adapted behaviors (Bond and Taylor, 2015).

As a final remark, our observation that adaptation is bounded by the synergistic space and that muscles and synergies are rotated only if engaged at their boundaries suggests a “greedy” adaptation process aiming at maximizing local efficiency (Emken et al., 2007; Ganesh et al., 2010), by which the association between muscular effort and workspace is modified only when necessary to the adaptation process, and left constant otherwise.

## Data Availability

The datasets generated for this study can be available on request to the corresponding author.

## Ethics Statement

The activities involving human participants were reviewed and approved by Ethic Committee, University College Dublin. The participants provided their written informed consent to participate in this study.

## Author Contributions

GS conceived the study, designed the experiments, analyzed the data and interpreted the results. GS and MZ performed the experiments and drafted the manuscript.

